# Optopharmacological tools for precise spatiotemporal control of oxytocin signaling in the central nervous system and periphery

**DOI:** 10.1101/2022.11.10.516001

**Authors:** Ismail A. Ahmed, Jing-Jing Liu, Krystyna A. Gieniec, Chloe J. Bair-Marshall, Ayomiposi B. Adewakun, Belinda E. Hetzler, Christopher J. Arp, Latika Khatri, Gilles C. Vanwalleghem, Alec T. Seidenberg, Pamela Cowin, Dirk Trauner, Moses V. Chao, Felicity M. Davis, Richard W. Tsien, Robert C. Froemke

## Abstract

Oxytocin is a neuropeptide critical for maternal physiology and social behavior, and is thought to be dysregulated in several neuropsychiatric disorders. Despite the biological and neurocognitive importance of oxytocin signaling, methods are lacking to activate oxytocin receptors with high spatiotemporal precision in the brain and peripheral mammalian tissues. Here we developed and validated caged analogs of oxytocin which are functionally inert until cage release is triggered by ultraviolet light. We examined how focal versus global oxytocin application affected oxytocin-driven Ca^2+^ wave propagation in mouse mammary tissue. We also validated the application of caged oxytocin in the hippocampus and auditory cortex with electrophysiological recordings *in vitro*, and demonstrated that oxytocin uncaging can accelerate the onset of mouse maternal behavior *in vivo*. Together, these results demonstrate that optopharmacological control of caged peptides is a robust tool with spatiotemporal precision for modulating neuropeptide signaling throughout the brain and body.

## Main

Oxytocin is an evolutionarily conserved neuropeptide with essential roles in many tissues across mammalian and many non-mammalian species (Jurek & Neumann, 2018; Gimpl & Fahrenholz, 2001; Theofanopoulou et al., 2021; Froemke & Young, 2021). In mammals, oxytocin is primarily synthesized in the paraventricular (PVN) and supraoptic nuclei (SON) of the hypothalamus (Althammer & Grinevich, 2017; Ludwig & Leng, 2006). The release of oxytocin into the bloodstream is critical for a number of peripheral physiological processes, including inducing uterine contractions during parturition to facilitate childbirth, and milk ejection during nursing (Valtcheva & Froemke, 2019; Numan & Young, 2016; Burbach et al., 2006). Oxytocin is also released throughout the central nervous system and has been shown to facilitate social behaviors such as maternal care (Marlin et al., 2015; Carcea et al., 2021) and pair bonding (Keebaugh et al., 2015; Walum & Young, 2018). Moreover, a deficit in oxytocin signaling is believed to be associated with several disorders, such that oxytocin supplements are being explored as a potential clinical treatment for obesity (Blevins & Baskin, 2015), pain (Mazzuca et al., 2011; Quintana, & Guastella, 2020), addiction (Kovács, et al., 1998), and psychiatric disorders such as autism spectrum disorders, social anxiety, posttraumatic stress disorder, and post-partum depression (Wagner & Harony-Nicolas, 2018; Bertsch & Herpertz, 2018; Frijling, 2017). However, the mechanisms of how oxytocin release and oxytocin receptor signaling modulate various cell types, tissues, and brain areas to regulate internal states and social interactions remains unclear. Thus far, the inability to control oxytocin peptide levels with high spatiotemporal precision *in vivo*, have hampered these efforts. For example, stimulating endogenous release or providing systemic oxytocin receptor agonists/antagonists to regulate signaling at specific sites or tissues without off-target effects has been challenging. Furthermore, both peptide and synthetic oxytocin receptor ligands can effectively bind and activate multiple receptor types present on the same cell or nearby cells. Direct peptide application *in vivo* and in brain slices by perfusion, pressure injection (Williams et al., 1982) or iontophoresis (Travagli et al., 1995) produces slow, prolonged and spatially imprecise presentation of the peptide offering poor control over the concentration and gradients of peptide delivered. Therefore, it is imperative to design new tools to enable spatiotemporally precise oxytocin delivery for regulating physiology and behavior.

In the periphery, circulating oxytocin is primarily released from the brain into the blood via the posterior pituitary, but there are also other peripheral sources of oxytocin (Paiva et al., 2021). These sources converge on their peripheral targets via diffusion (Chini et al., 2017). Oxytocin receptor expression is found in both neuronal and non-neuronal cell types in various organs including uterus, ovaries, testis, prostate gland, mammary, kidneys, heart, thymus, adipocytes, pancreas, adrenal glands, and others (Kimura et al., 2003; Gimpl & Fahrenholz, 2001). Consequently, oxytocin regulates a variety of critical physiological functions ranging from reproduction and lactation to control of body fluid and heart homeostasis (Kimura et al., 2003; Gimpl & Fahrenholz, 2001). Despite how fundamental oxytocin is in the periphery, spatiotemporal optogenetic studies of these different systems and processes have been inaccessible. Unlike in the brain, where the optogenetic release of oxytocin is, for the most part, practical (with some limitations and caveats), it is infeasible in the periphery. This is partly due at least to the lack of known oxytocin projections into peripheral tissues beyond the spinal cord and a lack of means to activate them (Eliava et al. 2016).

In the brain, previous studies of endogenous oxytocin release *in vivo* (e.g., through chemogenetic or optogenetic activation of hypothalamic oxytocin neurons) have provided essential information on the modulation of social behavior, mainly in rodents (Scott et al., 2015; Knobloch et al., 2012; Marlin et al., 2015). However, many hypothalamic cell types, including oxytocin neurons, are believed to co-express and release other neurotransmitters and peptides. Oxytocin-releasing neurons have been found to synthesize corticotropin-releasing hormone, cholecystokinin, and enkephalin (Martin & Voigt, 1981; Vanderhaeghen et al., 1981; Levin and Sawchenko, 1993), each of which has various functions for regulating physiological states and behavior (and could be released under different physiological conditions). Co-transmission of neuropeptides and small-molecules even by a single neuron can affect microcircuit activity and behavior through multiple contributing mechanisms and over a range of timescales (Nusbaum et al., 2017). Also, the spatial extent at which oxytocin signaling can affect a target brain region remains unclear. Oxytocin fibers into different brain areas do not seem to make conventional synapses. Fiber density may be sparse in many regions, and oxytocin can be released from the dendrites of hypothalamic neurons, suggesting that volume transmission must occur at least across some distances and in some regions (Mitre et al. 2016; Chini et al., 2017; Son et al., 2022). Additionally, there are many examples of projection—receptor mismatch across many brain regions which are mainly modulated by volume transmission and it may be difficult to activate with optogenetic approaches. Thus, compared to classical neurotransmitters, the spatial and functional specificity of oxytocin and other neuropeptides may be limited. Therefore, while genetics-based approaches have been useful for examining peptidergic modulation of neural circuit function, it can be difficult to parse out the contributions of specific co-transmitters from a particular cell population or the action of oxytocin in specific spatial locations or time points. Moreover, these approaches might be difficult to adopt for other species in which genetic manipulation is more challenging.

Here we developed optopharmacological compounds, variants of caged oxytocin, to precisely establish how oxytocin release drives receptor signaling in peripheral (in mouse mammary gland) and brain tissue (in mouse hippocampus and cortex) and to ultimately control maternal behavior. Optopharmacology enables a light-meditated selective activation of oxytocin signaling within a defined target region (Kramer et al., 2013; Hüll et al., 2018; Paoletti et al. 2019). This method complements optogenetics and pharmacology by endowing light sensitivity to specific biomolecular targets. It thus provides molecular control of a given system, in this case oxytocin receptor activation and regulation of mammary gland function in the periphery and neuronal activity in the central nervous system. This tool does not require genetic manipulations or viral expression and thus can be used in any organism that expresses the oxytocin receptor. Furthermore, our approach exemplifies that this technique can be generalized to small signaling peptides that target G-protein-coupled receptors (GPCRs) in the brain and body and would be applicable to both peptide agonists and antagonists for spatiotemporal specificity.

## Results

### Design and photophysical characterization of caged oxytocin analogs

Oxytocin is an ideal candidate for caging because of its small size (nine amino acids) and established role in various biological functions. Photosensitive caging has previously been successfully achieved for other peptides of similar size (Banghart et al. 2012; Montnach et al. 2022). To generate oxytocin that is functionally inert until light exposure, we produced analogs where individual amino acids were replaced with photo-caged unnatural amino acid versions of the corresponding residue using solid-phase peptide synthesis. Previous functional studies of oxytocin have shown that the Cys and Tyr residues are important for receptor activation (Barberis, et al., 1998; Donaldson et al., 2008). Additionally, the disulfide between the two Cys residues is responsible for the cyclic structure of oxytocin, which is important for the potency and metabolic half-life. Therefore, we hypothesized using photocage moieties on these residues would render oxytocin reversibly inert. Based on these considerations, we designed three caged oxytocin peptide analogs (**Fig. 1a**). For the first caged compound (‘cOT1’), we replaced Tyr with ortho-nitrobenzyl-tyrosine. The second analog (‘cOT2’) incorporated a 4,5-dimethoxy-2-nitrobenzyl caging group at the N-terminus of Cys^1^. For the third caged oxytocin compound (‘cOT3’), we replaced both cysteines with 4,5-dimethoxy-2-nitrobenzyl cysteine to impede cyclization through disulfide formation.

**Figure 1.**
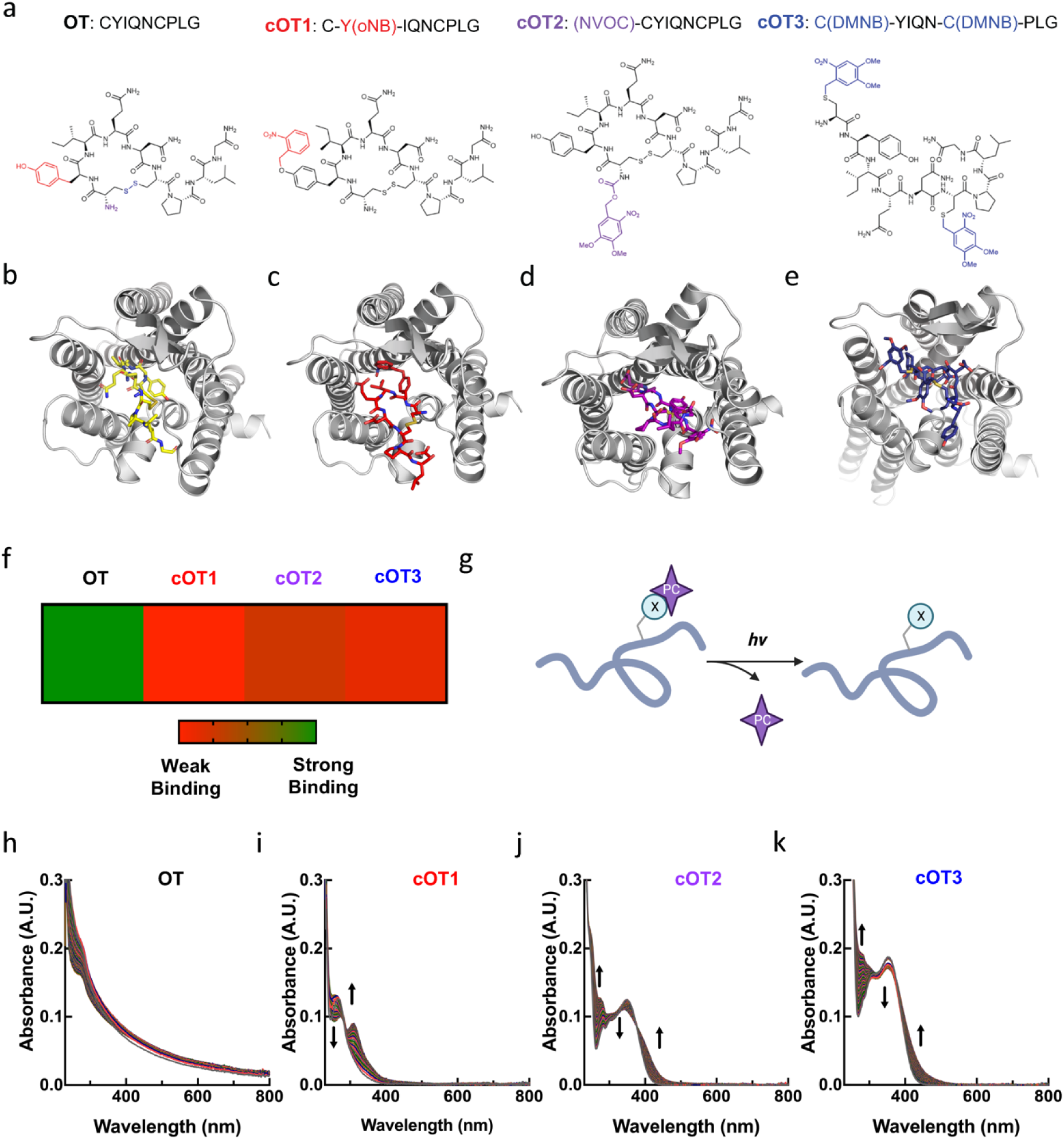
Structure, molecular modeling, and photophysical properties of oxytocin and caged oxytocin derivatives. (**a**) Amino acid sequences and chemical structures of oxytocin (OT) and the photocage-modified oxytocin analogs cOT1, cOT2, and cOT3. For the OT structure the functional groups that are modified with cages are color-coded: the Tyr residue is labeled in red, the N-terminus in purple, and the disulfide in blue. (**b**) Structure of OT bound to the oxytocin receptor, adapted from PDB: 7RYC. (**c-d**) Structure of photocaged-modified oxytocin analogs cOT1 (**c**), cOT2 (**d**), and cOT3 (**e**) bound to the oxytocin receptor (PDB: 7RYC) through molecular docking with Autodock Vina. (**g**) Heat map analysis of OT, cOT1, cOT2, and cOT3 screened against the oxytocin receptor (PDB: 7RYC) using Autodock Vina. In the gradient ruler, green color indicated strong binding, while red color indicates weak binding. Results listed from weakest binder: cOT1, cOT3, and cOT2. (**g**) General reaction scheme of uncaging with UV light to yield oxytocin from caged oxytocin; X: caged residue and PC: photocage. (**h-k**) Changes in the absorption spectrum of OT (**h**), cOT1 (**i**), cOT2 (**j**), and cOT3 (**k**) upon irradiation with 365 nm light (20 µM, room temperature, 1% DMSO/PBS). The spectra of each irradiated caged analog (cOT1, cOT2, and cOT3) show a significant change in absorbance, corresponding to photo-uncaging and oxytocin release.

To predict if each photocaged oxytocin analog (cOT1, cOT2, and cOT3) would is inert we performed molecular simulations of each compound in comparison to oxytocin bound to the oxytocin receptor (OXTR), PDB: 7RYC using Autodock Vina (**Fig. 1b-e**) (Meyerowitz et al., 2022). Each docked compound showed significant steric clashes residues within the binding pocket of OXTR and the weakest binding analog was cOT1, and then cOT3, and cOT2 respectively (**Fig. 1f**). Indeed, this suggested photocaging oxytocin would modify its binding affinity is due to alterations in the peptide/protein interaction. Furthermore, these results are in agreement with structural studies of oxytocin bound to OXTR. For example, the cryo-EM structure (PDB: 7RYC) shows the tyrosine residue of oxytocin penetrates deeply into the binding pocket of the helical bundle to form strong hydrophobic and hydrogen bond interactions with OXTR residues within (Meyerowitz et al., 2022). Undoubtedly, adding a bulky caging group (ortho-nitrobenzyl group) with starkly different chemical properties at this position as in the case of cOT1 will diminish its affinity for OXTR.

Photolysis of the selected caging groups can produce bioactive uncaged products on a millisecond time scale (Klán et al., 2012). Under optimal conditions, photorelease can proceed with >95% yield (Miller et al., 1998; Klán et al., 2012). We designed these caged oxytocin analogs for photorelease with 365 nm light and confirmed that the ultraviolet (UV) and visible absorption spectra of each analog corresponded to the uncaging wavelength of 365 nm (**Fig. 1a,g**). The spectrum of each caged oxytocin analog irradiated with UV light over 30 minutes (365 nm LED, 3 mW) showed a significant change in absorbance corresponding to photo-uncaging and release of oxytocin with UV light while oxytocin remained unchanged (**Fig. 1h-k**). Thus, oxytocin itself is stable towards 365 nm irradiation for durations of at least 30 minutes which is substantially higher than durations of irradiation that would be used in experimental conditions.

### *In vitro* validation of caged oxytocin analogs

We next determined if the caged oxytocin analogs cOT1, cOT2, and/or cOT3 are inactive at oxytocin receptors prior to photolysis and then become active after photolysis. We compared the activation of oxytocin receptors after uncaging to of the effects of oxytocin ligand wash-on using an *in vitro* functional calcium flux fluorescence assay (**Fig. 2a**). Oxytocin receptors are believed to primarily couple to Gq/11, mobilizing intracellular calcium (Gimpl & Fahrenholz, 2001). Thus, this assay should be robust for sensing receptor activation. To demonstrate the utility of uncaging of cOT1 in cell culture, we carried out Ca^2+^ imaging of CHO-K1 cells stably expressing oxytocin receptors and loaded the cells with the calcium-sensitive dye FLUO-8. We first treated cells expressing oxytocin receptors with Hanks balanced salt solution (HBBS) buffer and imaged under these control conditions before (**Fig. 2b**, top) and after photolysis (**Fig. 2b**, bottom) by full-field illumination with 365 nm UV light. We did not observe any change in fluorescence after UV illumination. We then treated these cells with 1 µM of cOT1 and again used full-field illumination with 365 nm UV light, observing a substantial increase in FLUO-8 fluorescence after photolysis (**Fig. 2c**).

**Figure 2.**
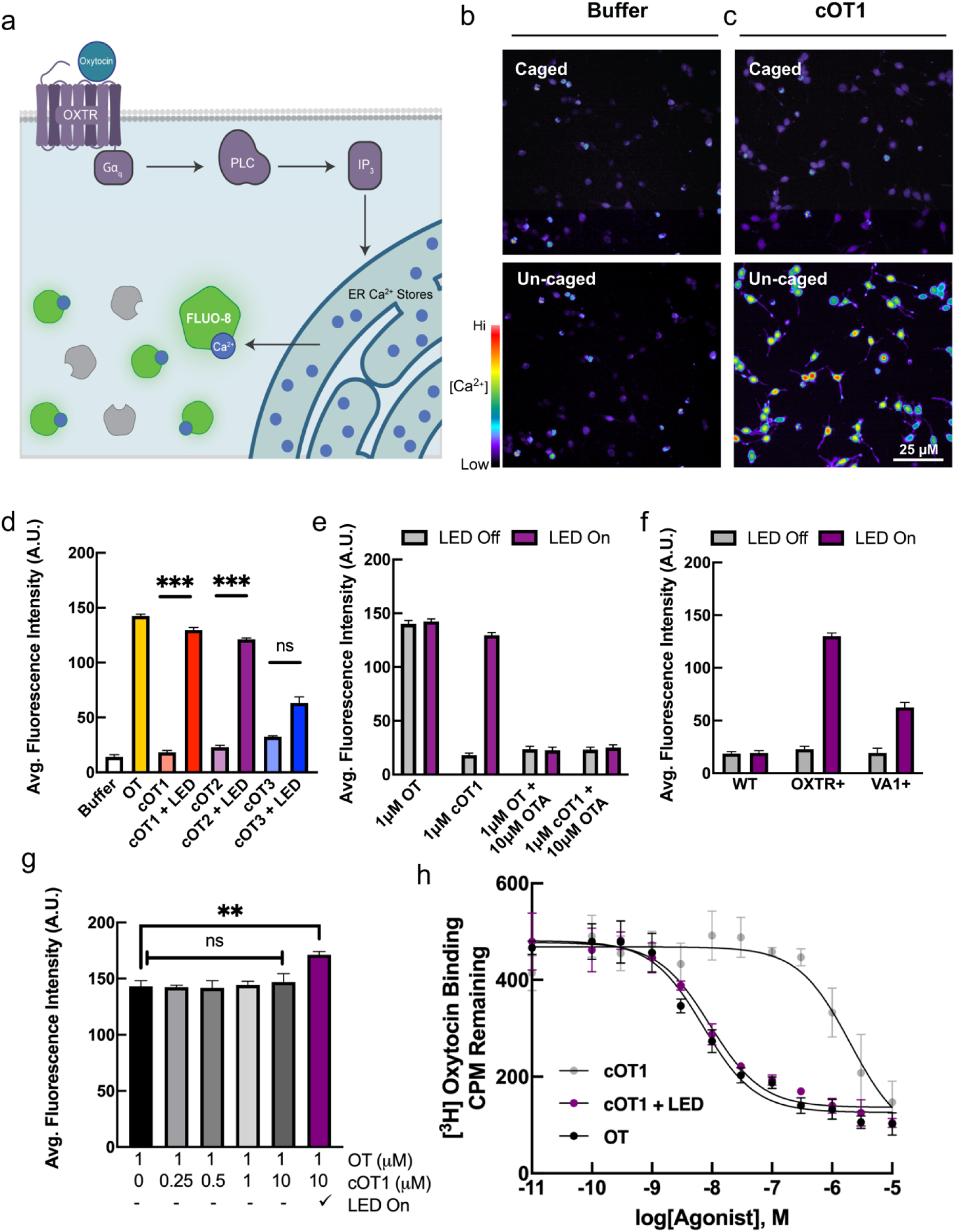
Caged oxytocin is inert before photolysis and activates the oxytocin receptor (OXTR) after photolysis with UV light. (**a**) Schematic of calcium flux fluorescence measurement of intracellular calcium release in CHO-K1 cells expressing OXTR and loaded with cell-permeable FLUO-8 calcium-sensitive dye. (**b**) Calcium imaging of CHO-K1 cells expressing OXTR loaded with FLUO-8 calcium-sensitive dye treated with HBBS buffer before (left) and after uncaging (right) with UV light (365 nm LED). (**c**) Calcium imaging of CHO-K1 cells expressing OXTR loaded with FLUO-8 calcium-sensitive dye treated with 1 µM of cOT1 before and after photolysis with UV light (365 nm LED). (**d**) Calcium flux assay to measure intracellular calcium release in CHO-K1 cells expressing OXTR and loaded with FLUO-8 calcium-sensitive dye treated with either buffer (HBBS), oxytocin (HBBS, fluorescence: 14.1±2.0, n=5; oxytocin, fluorescence: 142.4±1.8, n=5 cell culture wells, p=0.0004, Student’s two-tailed paired t-test), and the caged oxytocin analogs cOT1 (cOT1 LED off, fluorescence: 18.1±1.9, n=5 cell culture wells; cOT1 LED on, fluorescence: 129.7±2.4, n=5 cell culture wells, p=0.0007, Student’s two-tailed paired t-test), cOT2 (cOT2 LED off, fluorescence: 22.8±2.1, n=5 cell culture wells; cOT2 LED on, fluorescence: 121.1±1.4, n=5 cell culture wells, p=0.0008, Student’s two-tailed paired t-test), cOT3 (cOT3 LED off, fluorescence: 40.4±1.9, n=5 cell culture wells; cOT3 LED on, fluorescence: 85.2±3.3, n=5 cell culture wells, p=0.0008, Student’s two-tailed paired t-test) before and after photolysis with UV light (365 nm LED) at a concentration of 1 µM respectively. Data were normalized to the maximal response produced by oxytocin, the endogenous agonist of OXTR; error bars: mean ± SEM. (**e**) Calcium flux assay using CHO-K1 cells expressing OXTR treated with oxytocin or cOT1 with and without OTA, an inhibitor for OXTR before and after photolysis with UV light (365 nm LED) at a concentration of 1 µM and 1 µM respectively. Data were normalized to the maximal response produced by oxytocin; n= 5 cell culture wells per condition; for 1 μM OTA p=0.17 and 10 μM OTA p=0.28 error bars: mean ± SEM. (**f**) Calcium flux assay using wild type CHO-K1 cells and CHO-K1 cells expressing OXTR or vasopressin receptor (VA1) treated with cOT1 before and after photolysis (OXTR: cOT1 LED off, fluorescence: 22.7±2.8, n=5 cell culture wells; cOT1 LED on, fluorescence: 130.1±2.9, n=5 cell culture wells, p=0.0005; VA1: cOT1 LED off, fluorescence: 19.3±4.5, n=5 cell culture wells; cOT1 LED on, fluorescence: 62.4±4.8, n=5 cell culture wells, p=0.0003 cell culture wells; Student’s two-tailed paired t-test). Data were normalized to the maximal response produced by oxytocin; error bars: mean ± SEM. (**g**) Calcium flux assay using CHO-K1 cells expressing OXTR treated with oxytocin or cOT1 (at various concentrations) with and without 365 nm light to test cOT1 for antagonism of OXTR. Data were normalized to the maximal response produced by oxytocin; error bars: mean ± SEM. Cells treated with both 1 µM oxytocin and 10 µM cOT1 with LED on showed an increase in calcium signal compared to oxytocin alone (1 µM oxytocin 10 µM cOT1 LED on, fluorescence: 171.2±2.9, n=5 cell culture wells, p=0.0043, Student’s two-tailed paired t-test). The varying concentrations of cOT1 with oxytocin and LED off are n.s. when compared to oxytocin. (**h**) Competitive binding curves OXTR of cOT1 either caged (left) or uncaged (right) relative to the endogenous OXTR agonist, oxytocin. The affinity for OXTR by oxytocin (IC_50_ = 7.16 nM), caged cOT1 (IC_50_ = 2.01 mM), and uncaged cOT1 (IC_50_ = 9.24 nM), paired t-test. ** p<0.01.

**Extended Data Figure 1.**
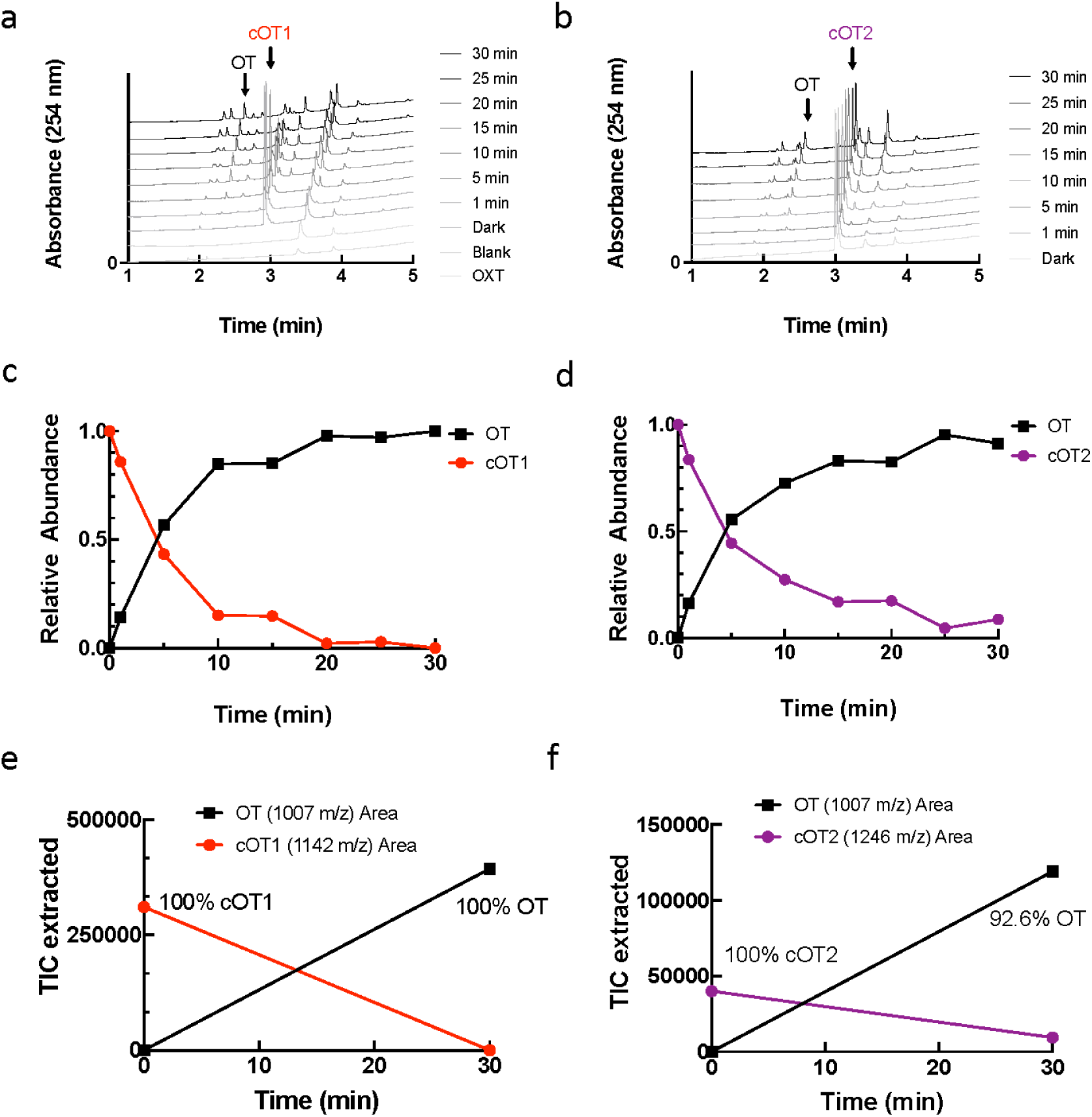
Analysis of Uncaging of cOT1 and cOT2 using LC-MS. (**a-b**) Representative set of LC-MS traces for cOT1 (**a**) and cOT2 (**b**) show an increase in oxytocin and simultaneous decrease in photocaged oxytocin upon 365 nm irradiation. LC-MS Data: OT: LC-MS t_R_ = 2.19 min (5–100% MeCN in H_2_O, 0.1% formic acid, 254 nm detection) LRMS (ESIP) for C H N O S ^+^ [M+H]^+^: calcd.: 1007.4, found 1007.4. cOT1: LC-MS t_R_ = 2.82 min (5–100% MeCN in H_2_O, 0.1% formic acid, 254 nm detection) LRMS (ESIP) for C H N O S ^+^ [M+H]^+^: calcd.: 1142.48, found 1142.5. cOT2: LC-MS t_R_ = 2.97 min (5–100% MeCN in H_2_O, 0.1% formic acid, 254 nm detection) LRMS (ESIP) for C H N O S ^+^ [M+H]^+^: calcd.: 1246.5, found 1246.5. (**c-d**) Plot of photouncaging of cOT1 (**c**) and cOT2 (**d**). (**e-f**) After 20 min full uncaging of cOT1 (**e**) is observed. After 30 min, 92.6% of cOT1 (**f**) uncaging observed.

In total, we carried out *in vitro* functional calcium flux fluorescence assay from CHO-K1 cells stably expressing oxytocin receptors and loaded with the calcium-sensitive dye FLUO-8, that were treated with HBBS buffer, oxytocin (1 µM), or one of the caged oxytocin analogs (cOT1, cOT2, cOT3; 1 µM) before and after photolysis with UV light with an LED (365 nm, 3 mW). Before photolysis, cOT1 and cOT2 demonstrated minimal activation of oxytocin receptors as inferred from calcium imaging, comparable to that measured in HBBS buffer but cOT3 showed significant activation (**Fig. 2d**; HBBS, fluorescence: 14.1±2.0, n=5 cell culture wells; cOT1 LED off, fluorescence: 18.1±1.9, n=5 cell culture wells, p=0.23; cOT2 LED off, fluorescence: 22.8±2.1, n=5 cell culture wells, p=0.16; cOT3 LED off, fluorescence: 40.4±1.9, n=5 cell culture wells, p=0.0008; Student’s two-tailed paired t-test). After photolysis, both cOT1 and cOT2 produced robust calcium signals, similar to that produced by oxytocin itself when compared to HBBS (**Fig. 2d**; HBBS, fluorescence: 14.1±2.0, n=5 cell culture wells; oxytocin, fluorescence: 142.4±1.8, n=5 cell culture wells, p=0.0004; cOT1 LED off, fluorescence: 18.1±1.9, n=5 cell culture wells; cOT1 LED on, fluorescence: 129.7±2.42, n=5 cell culture wells, p=0.0007; cOT2 LED off, fluorescence: 22.8±2.1, n=5 cell culture wells; cOT2 LED on, fluorescence: 121.1±1.4, n=5 cell culture wells, p=0.0008, Student’s two-tailed paired t-test). To compare the photo-release of pure oxytocin from cOT1 and cOT2, we used standard liquid chromatography–mass spectrometry and we observed full photo-uncaging of cOT1 after 20 min and ~90% uncaging is observed for cOT2 after 30 minutes (**Extended Data Fig. 1**). These experiments confirm that the caged oxytocin analogs we synthesized can effectively photorelease oxytocin peptides. For the remainder of this study, we only used cOT1 because this reagent was the most inert of the caged-oxytocin analogs before photolysis and showed the most robust activation of oxytocin receptors after photolysis.

To demonstrate that the effects of cOT1 uncaging are specific for the oxytocin receptor, we performed similar UV photolysis experiments with cOT1 onto CHO-K1 cells expressing oxytocin receptors, and included the specific oxytocin receptor antagonist OTA in the bath. When inhibiting OXTR with OTA, cOT1 neither activates OXTR cells before or after photolysis, thus demonstrating the uncaging of oxytocin (cOT1) is specific to OXTR binding (**Fig. 2e**, for 1 μM OTA p=0.17, 10 μM OTA p=0.28). Furthermore, uncaging cOT1 in the presence of VA1 vasopressin receptor expressing cells shows less activation in comparison to uncaging cOT1 in the presence of OXTR expressing cells (**Fig. 2f**; OXTR: cOT1 LED off, fluorescence: 22.7±2.8, n=5 cell culture wells; cOT1 LED on, fluorescence: 130.1±2.9, n=5 cell culture wells, p=0.0005; VA1: cOT1 LED off, fluorescence: 19.3±4.5, n=5 cell culture wells; cOT1 LED on, fluorescence: 62.4±4.8, n=5 cell culture wells, p=0.0003; Student’s two-tailed paired t-test), as we would expect from the difference in affinity of oxytocin for the cognate receptor OXTR versus the V1A receptor.

To ensure that cOT1 had no detectable antagonism to OXTR in experimental concentration ranges, we carried out the same calcium flux assay described above. Increasing concentrations of cOT1 (0 µM, 0.25 µM, 0.5 µM, 1 µM, 10 µM) were applied in bath to CHO-K1 cells stably expressing oxytocin receptors loaded with the calcium-sensitive dye FLUO-8 and then subsequently treated with 1 µM of oxytocin and no UV light (LED off). Over the entire concentration range, there was no significant change (e.g., reduction) in the calcium signal when compared to cells treated solely with 1 µM oxytocin (**Fig. 2g**). However, when cells were treated with both 1 µM oxytocin and 10 µM cOT1 with LED on, there was an expected increase in calcium signal compared to oxytocin alone (**Fig. 2g**; 1 µM oxytocin LED off, fluorescence: 143.3.1±4.9, n=5 cell culture wells; 1 µM oxytocin 10 µM cOT1 LED on, fluorescence: 171.2±2.9, n=5 cell culture wells, p=0.0043, Student’s two-tailed paired t-test). These data suggest that cOT1 is not antagonistic, and thus this compound might enable uncaging experiments in which balance between activation and inhibition in a receptor is important, such as in *ex vivo* tissue and in freely behaving animals.

To quantify the difference in binding affinity of cOT1 compared to oxytocin, we obtained dose-dependent competitive binding curves of cOT1 and oxytocin, either caged or uncaged (**Fig. 2h**). The affinity of cOT1 (IC_50_ = 2.01 µM) is ~280 times less than oxytocin (IC_50_ = 7.16 nM). After photolysis, the affinity of cOT1 for OXTR is similar to that of oxytocin (IC_50_ = 9.24 nM). These data reveal caging at the Tyr residue of oxytocin has the most robust effect on its binding and activation of OXTR.

### Photorelease of oxytocin in mammary tissue

Oxytocin is critically important for the milk ejection reflex; knockout mice lacking either oxytocin peptide or oxytocin receptors are unable to expel milk to support their young (Nishimori et al., 1996; Takayanagi et al., 2005). With other methods such as systemic release or optogenetic stimulation of the hypothalamus, precise regulation of mammary function can be especially challenging because oxytocin is released into the bloodstream from the posterior lobe of the pituitary gland. Thus, we next sought to demonstrate the utility of caged oxytocin in mammary tissue. During lactation, mammary glands express high levels of oxytocin receptors (**Fig. 3a,b**). We performed three-dimensional time-lapse imaging of *ex vivo* mammary tissue from lactating GCaMP6f::K5CreERT2 mice treated with cOT1 and subsequently bathed with oxytocin. With a 405 nm laser, we uncaged a region of interest (ROI) at 100% laser power for 3 seconds at three different time points. Uncaging of cOT1 only occurred in the radius of the ROI, as measured by increases in cytosolic calcium and alveolar movement, but did not occur in cells distal from the ROI (**Fig. 3c,d**). However, as previously shown (Stevenson et al., 2020), the star-shaped basal epithelial cells robustly respond to oxytocin with an increase in intracellular calcium, followed by cell and alveolar contraction across all cells throughout the sample in a diffusive manner after bath application (**Fig. 3c,d**). We observed that uncaging on one alveolar cluster typically led to local activation just of that cluster, and did not seem to induce global synchrony of calcium release and contractions throughout adjacent mammary tissue. This implies that global alveolar contraction and milk ejection requires a minimum concentration and more diffuse spatial distribution of oxytocin in the tissue.

**Figure 3.**
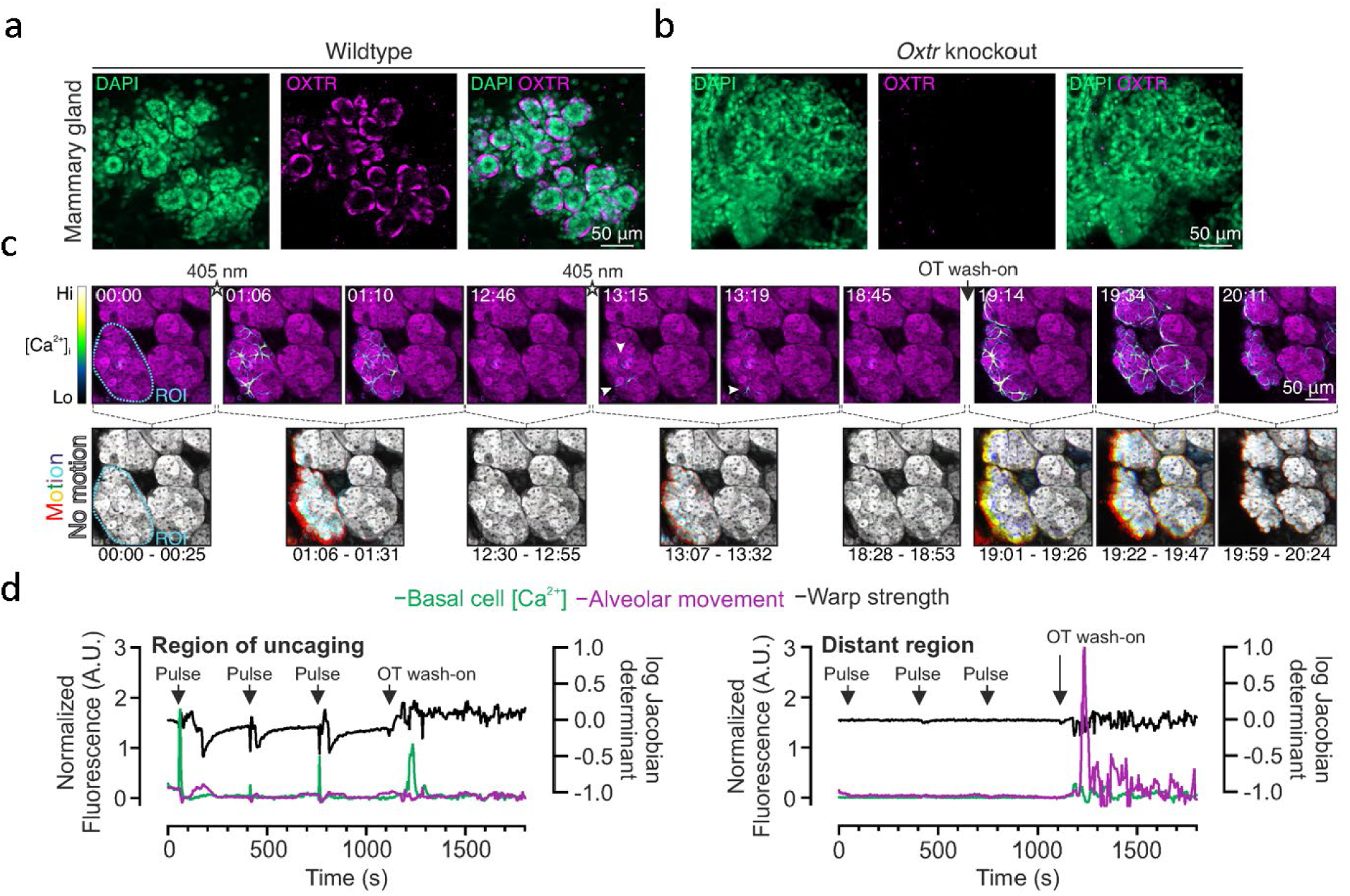
Photorelease of oxytocin in mammary tissue induces basal cell Ca^2+^ oscillations and alveolar contractions. (**a**) Image of cleared lactating mammary tissue of WT mouse, showing immunolabeling of oxytocin receptors with OXTR-2 antibodies (magenta). Nucleus labeled with DAPI (green). (**b**) Image of cleared lactating mammary tissue of OXTR KO mouse. There was no obvious OXTR-2 staining in the KO tissue. (**c**) Three-dimensional time-lapse imaging of live mammary tissue from GCaMP6f::K5CreERT2 lactating mice loaded with CellTracker Red (magenta) and incubated with cOT1 (85 nM) for 5 minutes and then uncaged cOT1 in an ROI with 405 nm light at 100% laser power for 3 seconds (3 trials) and subsequently bathed with 85 nM of oxytocin. Top panels: Calcium signal; Bottom panels: Alveolar movement analysis generated by the overlay of three images (25 s apart). Each image has been assigned a primary color. Regions that do not move during the window have R-G-B pixels superimposed and are white. Regions where significant movement has occurred appear R, G, B, or a combination of two colors. (**d**) Overlay of basal cell calcium signal, alveolar movement (left axis), and warp strength (right axis) of region of uncaging (left panel) versus a distant region away from uncaging ROI radius (right panel).

### Photorelease of oxytocin in neural tissue

To validate the ability to uncage cOT1) (**Fig. 4a**) to activate OXTR^+^ neurons in the brain, we first sought to carry out a functional assay on the depolarizing effect of oxytocin previously identified in OXTR^+^ neurons in the hippocampus (Tirko et al., 2018; Liu et al., 2022) OXTRs are highly expressed in CA2 pyramidal neurons in the hippocampus (Mitre et al., 2016), and their activation drives neuronal depolarization (Tirko et al., 2018; Liu et al., 2022). We visually targeted OXTR-expressing (OXTR^+^) neurons in the dorsal CA2 region (dCA2) (**Fig. 4b**), using OXTR-Cre mice crossed with an Ai9 tdTomato reporter line (Dudek et al., 2016; Raam et al., 2017; Young et al., 2020).

**Figure 4.**
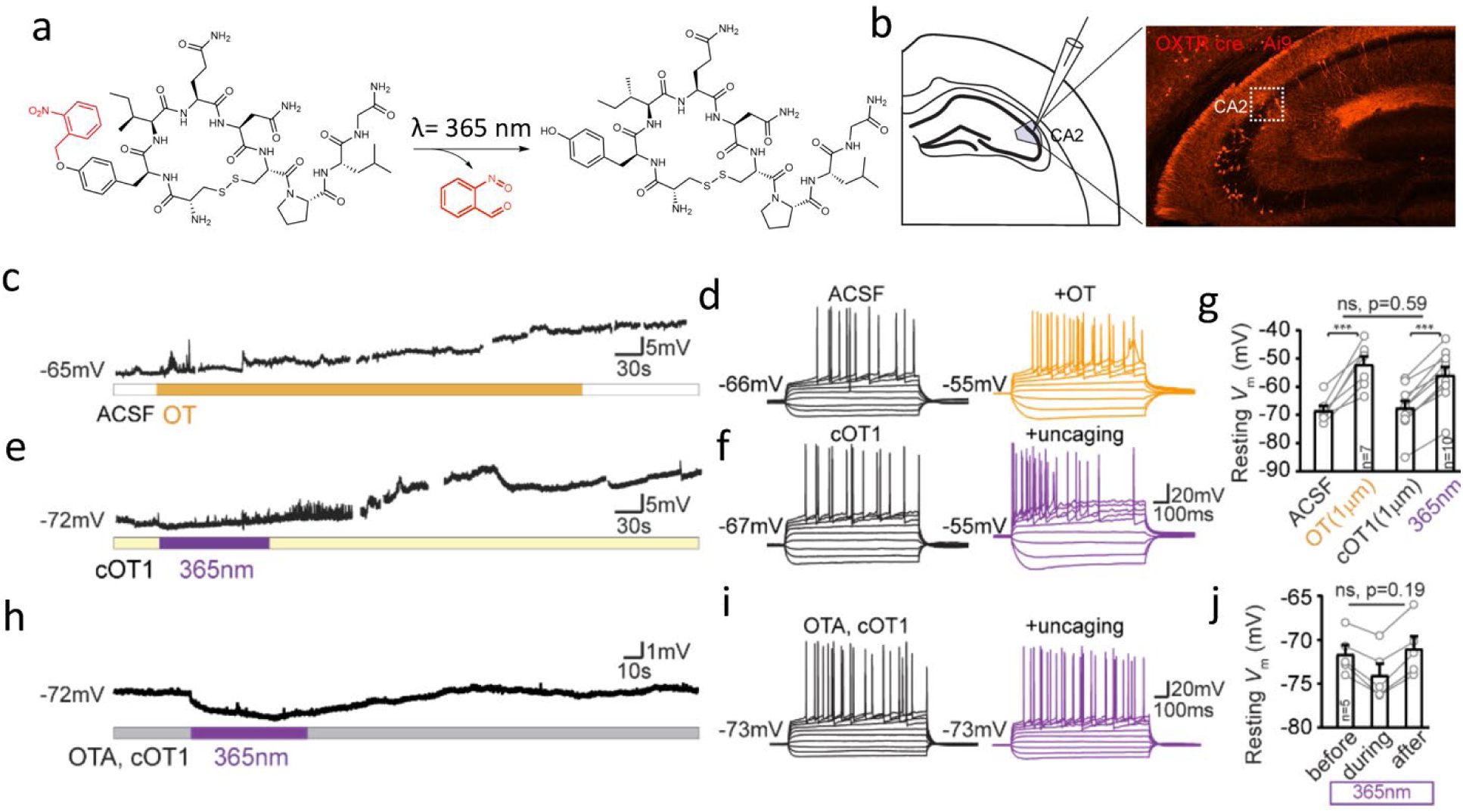
Photo-release of caged oxytocin depolarizes OXTR^+^ hippocampal CA2 pyramidal neurons. (**a**) Uncaging reaction of cOT1. Exciting the chromophore of ortho-nitrobenzyl-tyrosine with UV light should yield as photoproducts primarily oxytocin and 2-nitrosobenzaldehyde. (**b**) Schematic of whole-cell recordings from mouse hippocampal brain slices (left), image of a coronal section containing dorsal hippocampus CA2 from OXTRcre::Ai9 mouse where neurons expressing oxytocin receptors express tdTomato (right). (**c**) Example traces of spontaneous activity recorded on current clamp of tdTomato^+^ pyramidal cell during application of exogenous 1 µm oxytocin (orange bar). (**d**) Example traces of firing patterns of CA2 OXTR^+^ pyramidal cell induced by steps of current injection (−200Δ40pA) before (black) and after (orange) application of 1 µm oxytocin (**e**) Example traces of spontaneous activity of tdTomato^+^ dCA2 neuron in the presence of 1 µm cOT1 and UV light stimulation (purple bar). (**f**) Example traces of firing patterns before (black) and after UV photo-uncaging of cOT1 (purple). (**g**) Pooled data of resting membrane potential (*V*_m_) quantified at the 15^th^ minute since oxytocin in or light-on respectively. Gray dots represent individual cells. Bar graphs represent Mean ± SEM: in ACSF application of oxytocin depolarized CA2 neurons from −68.43±1.59 to −52.14±2.89, n=7, p=0.0009, paired t-test; UV light-mediated uncaging depolarized CA2 neurons from −67.55±2.53 to −55.89±2.90, n=10, p<0.0001, paired t-test. Photo-uncaging depolarized CA2 neurons to a similar extent as did oxytocin, F (1, 23) = 0.2962, p=0.5915, Two-way ANOVA. (**h**) Example traces of spontaneous activity recorded on current clamp of tdTomato+ pyramidal cell in bath solution containing 10 µm OTA and 1 µm cOT1, and photo-stimulation was applied using 365 nm UV light (purple bar). (**i**) Firing patterns of CA2 cell in the presence of OTA and cOT1, before (black) and after (purple) 365 nm UV light stimulation. (**j**) Pooled data of resting membrane potential (*V*_m_). Gray dots represent individual cells. Bar graphs represent Mean ± SEM: before, during and after 365 nm LED light stimulation the *V*_m_ was −71.6±1.04, −74.17±1.15, and −71.02±1.43 mV, F (2, 12) = 1.900, p=0.1919, One-way ANOVA. ** p<0.01, ***p<0.001

Bath application of 1 µM exogenous oxytocin strongly depolarized the membrane potential (*V*_m_) of dCA2 OXTR^+^ neurons (from −68.4±1.6 to −52.1±2.9 mV, n=7, p=0.0009) (**Fig. 4c**) and altered their action potential firing pattern (**Fig. 4d**) as previously shown (Tirko et al., 2018). In comparison, bath perfusion of caged cOT1 at 1 µM without UV uncaging induced no significant change in CA2 pyramidal cells (68.4±1.6, n=7 vs. −67.6±2.5 mV, n=10, p=0.80, unpaired t-test) (Fig. 4d,e). However, additional 1-2 minutes of 365 nm UV light induced changes in firing pattern as well as resting *V*_m_ (from −67.6±2.5 mV to −55.9±2.9 mV, n=10 neurons, p<0.0001, unpaired t-test), to a similar degree as 1 µM oxytocin (two-way ANOVA, p=0.59) (**Fig. 4c-g**). We confirmed that the effects of cOT1 uncaging were due to OXTR activation, as both the firing pattern and the depolarization driven by UV uncaging were blocked by pre-application of 10 µM OTA, a specific OXTR antagonist (**Fig. 4h-j**).

We also asked oxytocin uncaging would have similar effects on neurons in another brain region, in this case the mouse auditory cortex. We have previously reported lateralized OXTR expression to the left auditory cortex (AC), and the increased OXTR expression has been shown to be important for maternal behaviors such as retrieving isolated pups back to the nest (Marlin et al., 2015; Mitre et al., 2016; Carcea et al., 2021). We targeted our recordings to OXTR-expressing (OXTR+) neurons in the auditory cortex (**Fig. 5a**) using OXTRcre::Ai9 mice (Dudek et al., 2016; Raam et al., 2017; Young et al., 2020). We performed whole-cell recordings in acute brain slices of the mouse auditory cortex. We found that photorelease of the caged oxytocin caused significant membrane depolarization of OXTR+ neurons (n=9 neurons, p =0.0003, paired t-test) (**Fig. 5b,c**). The effect of photolyzed cOT1 was similar to the depolarization produced by pharmacological application of oxytocin (**Fig. 5b,d**; n=10 neurons, p =0.0013, paired t-test). Together, these *in vitro* experiments in CA2 and auditory cortex demonstrate that cOT1 is a robust optopharmacological tool to activate OXTRs in neuronal tissue. Additionally, this compound can be applied to slices for hours without apparent toxic effects on neurons. We suggest that this tool will enable spatiotemporal delivery of oxytocin directly to the site of action over a range of areas and photorelease oxytocin in an analog or gradient manner by varying the amount of photolysis light (power and time) as demonstrated in previous studies with caged compounds (Banghart et al. 2012; Durand-de Cuttoli et al. 2020; Montnach et al. 2022).

**Figure 5.**
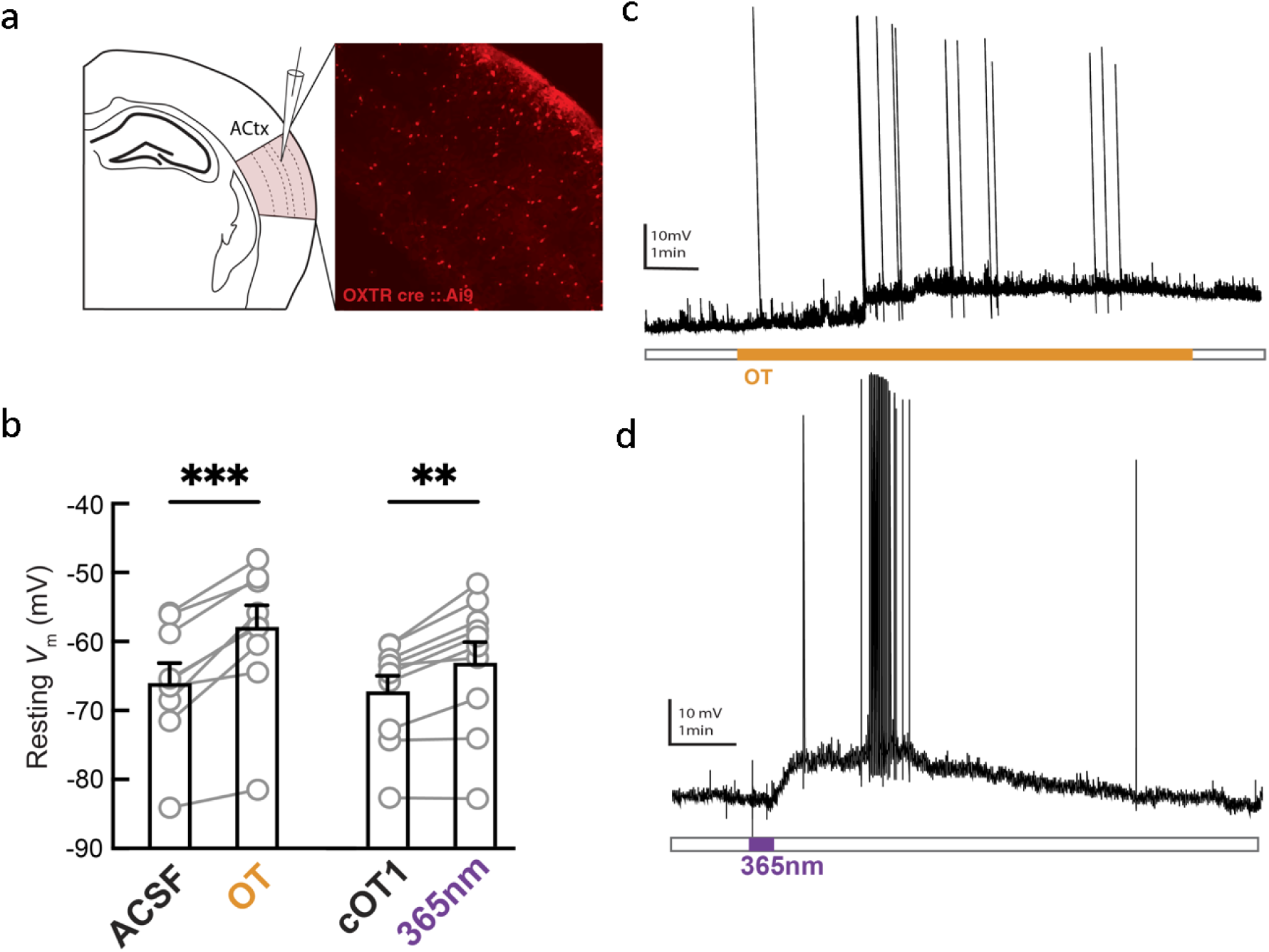
Photorelease of oxytocin depolarizes OXTR+ neurons in the auditory cortex. (**a**) Schematic of slice recordings (right), image of a section of left auditory cortex from OXTRcre::Ai9 mouse where neurons expressing oxytocin receptors express tdTomato (left). (**b**) Summary graph of peak depolarization of OXTR+ neurons for oxytocin (left) and caged oxytocin (cOT1) (right). Photorelease of caged oxytocin (cOT1) causes significant membrane depolarization of OXTR+ neurons (n=10 neurons, p =0.0013, paired t-test) and compares well (n.s. difference, unpaired t-test) with the pharmacological application of oxytocin which produces significant membrane depolarization of OXTR+ neurons (p = 0.0003, paired t-test). (**c**) Example trace of continuous current clamp recording from OXTR+ neuron (red) and OXTR-neuron (black) with oxytocin peptide. Example trace of continuous current clamp recording from OXTR+ neuron (black) and OXTR-neuron (grey) with caged oxytocin (cOT1), pulsed with UV light (365 nm LED) for uncaging at the purple arrow. (**d**) Example trace of continuous current clamp recording from OXTR+ neuron (black) and OXTR-neuron (grey) with caged oxytocin (cOT1), pulsed with UV light (365 nm LED) for uncaging at the purple arrow. ** p<0.01, ***p<0.001

### Photorelease of oxytocin enables control of maternal behavior

Finally, we asked if caged oxytocin could be useful *in vivo* for regulation of behavior; in this case, the emergence of mouse alloparenting behavior. One important mouse maternal behavior is pup retrieval; when pups are separated from the nest, they make ultrasonic distress calls, which experienced mothers (dams) then use as a cue to retrieve the isolated pups. Initially, most inexperienced (pup-naïve) virgin female mice do not retrieve pups but can become alloparents during co-housing with dams. This process requires native oxytocin and is accelerated by oxytocin supplements (Marlin et al. 2015; Schiavo et al., 2020; Carcea et al., 2021).

To deliver cOT1 into the mouse auditory cortex *in vivo*, we implanted an optofluidic cannula positioned in the left auditory cortex of pup-naïve female virgin C57Bl/6 mice (**Fig. 6a**), and after each experiment we verified the position of the cannula (**Fig. 6b**). Virgin females were co-housed with experienced dams and their litters over several days, and we initially verified that dams retrieved pups (**Fig. 6c-e**). Two cohorts of virgin females were infused with cOT1 and were tested for pup retrieval (**Fig. 6d**); one experimental ‘LED on’ group for uncaging cOT1, and one control ‘LED off’ to retain cOT1 as functionally inert (**Fig. 6e-f**). Within 12 hours of co-housing with a dam and her pups, virgin females with the LED on after cOT1 infusion began retrieving more than the LED off cohort (**Fig. 6e**; off ‘LED on’: 7 / 9 animals retrieved; LED off: 3 / 12 animals retrieved, p=0.03, two-tailed Fisher’s exact test). This meant that the emergence of this alloparental behavior was slower for the LED off cohort compared to the LED on cohort, although eventually most animals in both groups began retrieving (**Fig. 6f**), as expected from past work on co-housing (Marlin et al. 2015; Schiavo et al., 2020; Carcea et al., 2021).

**Figure 6.**
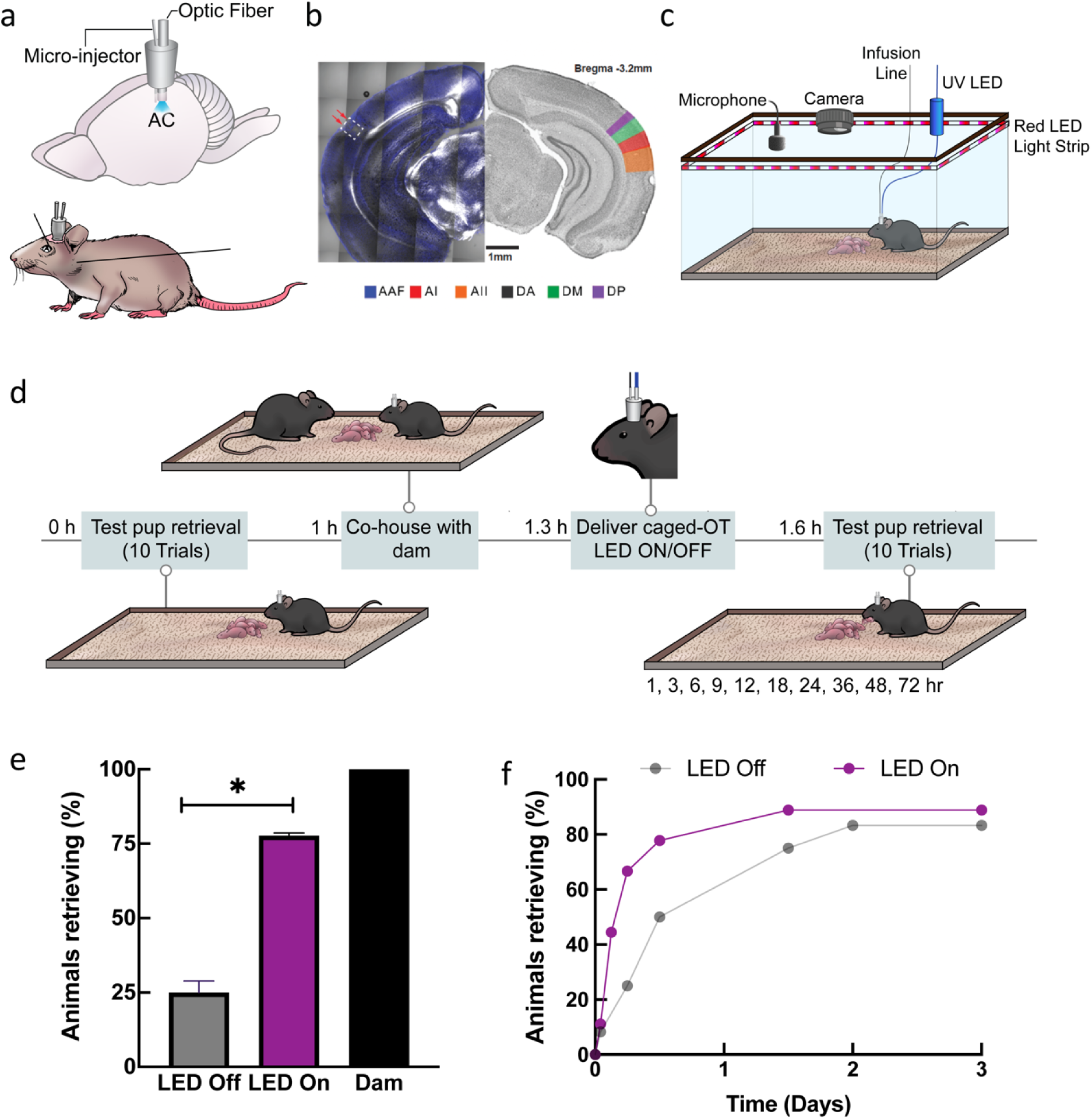
Photorelease of oxytocin in the auditory cortex enables pup retrieval in virgin mice. (**a**) Cartoon of an implanted optofluidic cannula to deliver both light and fluid in the mouse brain. (**b**) Histology validating implantation of optofluidic cannula targeting the left auditory cortex. (**c**) System for monitoring pup retrieval behavior compatible with an optofluidic cannula. (**d**) Schematic of pup retrieval behavior paradigm of virgin female mice co-housed with experienced dams comparing caged (LED Off) and uncaged (LED On) oxytocin. (**e**) Initially-naive virgins reliably retrieving 2+ times <12 hours after co-housing while infused with caged oxytocin with either with the 365 nm LED on or off (LED On: 7 / 9 animals; LED Off: 3 / 12 animals), p=0.03 was determined using Fisher’s exact test (two-tailed); Error bars: means ±95% confidence intervals. (**f**) Cumulative retrieval during co-housing. * p<0.05

## Discussion

In this study, we designed several photoactivatable oxytocin analogs by replacing key residues (Cys and Tyr) with their photocaged unnatural amino acid complement. We characterized the photophysical and photochemical properties of each caged oxytocin peptide to ensure successful uncaging with UV light. We compared each caged oxytocin analog using calcium sensitive assays in cell culture as a proxy for the activation of OXTR in the absence and presence of UV light. cOT1 was caged at the Tyr residue hydroxyl group, and we found this compound to be the best performing of the caged oxytocin derivatives, without antagonistic effects on the oxytocin receptor under our experimental conditions. The binding affinity of cOT1 to the oxytocin receptor in the caged form is ~280X less than in the uncaged form, where after photolysis with UV light, its binding affinity for the oxytocin receptor is approximately the same as oxytocin (because the reaction yields native oxytocin). These results point to the Tyr residue of oxytocin as the most effective caging site to attenuate the potency of. In this study, we specifically used the ortho-nitrobenzyl photo labile group to cage cOT1, which has been successfully used in various biological applications, including the photocontrol of proteins such as nicotinic acetylcholine receptors (Miller et al., 1998), protease activity (Luo et al., 2017), and has been established as a general tool for turning off and on protein activity (applied to GTPases, kinases, RNA demethylases, caspases and bacterial effector proteins) through caging and decaging actions (Wang et al., 2019). Other variations of caged tyrosine unnatural amino acids analogs have also been successfully used (Luo et al., 2017). The design of cOT1 serves as a scaffold for designing and tuning the photoactivation properties of caged oxytocin. For example, the caging group on the Tyr can be replaced with other caging functional groups with distinct chemical and physical properties such as to optimize for faster photolysis or multiphoton uncaging. Together, this study serves as a model for designing photoactivatable derivatives of similar neuropeptides of similar sizes, such as the closely related neuropeptide, vasopressin and can further be applied to endow photocontrol of specific peptidergic agonists and antagonists.

Oxytocin signaling is fundamentally important in non-neuronal tissues, most notably the mammary gland where oxytocin receptor activation triggers milk ejection in response to infant suckling (Valtcheva & Froemke, 2019). Basal epithelial cells respond to oxytocin with increased intracellular calcium, followed by cell and alveolar contraction (Stevenson et al., 2020). Both stochastic and coordinated events have been observed in mammary tissue, and mechanisms for cell entrainment and tissue-level synchronization are the topic of ongoing investigations. Caged compounds may be particularly advantageous for deciphering cellular connectivity in the mammary gland by optical spatiotemporal activation, particularly since genetic methods present challenges in this epithelial system. We show that caged oxytocin can be released in thick, lactating mammary tissue with a brief 405 nm pulse, causing local release of oxytocin and alveolar unit contraction. These findings point to the utility of this synthetic compound, both with conventional microscopes and in non-neuronal systems.

As the toolbox of optical tools in biology continues to grow, combining these tools becomes an increasingly powerful approach for dissecting information within brain circuits and other organs. Combining these tools will allow for the control and monitoring of more signals simultaneously. For example, a useful feature of using caged oxytocin is that it allows for three-channel (potentially more) multiplexing for imaging studies, as we have shown with the imaging of contracting mammary tissue activated by uncaging oxytocin (**Fig. 3**). Specifically, uncaging oxytocin with 405 nm light (or 365 nm), and then online imaging of calcium dynamics using GCaMP6f and cell structural changes using CellTracker Red in the green and red channels correspondingly. This strategy has the potential to be useful in cases where oxytocin can be uncaged and have two different readouts in the green and red channels with a combination of sensors and/or structural markers. Additionally, uncaging oxytocin can be used in parallel with optogenetic stimulation and inhibition because of minimal spectral overlap in the excitation wavelength between the tools. In the oxytocin system, there are several studies that show how oxytocin signaling affects modulates fast neurotransmitter action, for example, oxytocin modulation of dopamine neurons in the VTA (Xiao, et al., 2018), and there is evidence that the oxytocin receptor can form heterocomplexes with other receptors like the dopamine and serotonin receptors (Romero-Fernandez et al., 2013; Chruścicka et al., 2021). The design of this caged oxytocin opens up opportunities for manipulation of the oxytocin signaling without the use of genetics, thus enabling spatiotemporal control without having to use genetic manipulation or viral expression of opto/chemogenetic tools. Moreover, caged-oxytocin has the potential to be a useful tool for optical control of oxytocin binding and signaling in non-model organisms where genetic methods are not yet available.

## Methods

All procedures carried out at NYU Grossman School of Medicine were approved under NYU Grossman School of Medicine IACUC protocols. Animal experimentation done at the University of New South Wales (Sydney, Australia) was in accordance with the Australian Code for the Care and Use of Animals for Scientific Purposes with local animal ethics committee approval.

### Synthesis

All caged oxytocin analogs (cOT1, cOT2, and cOT3) were synthesized using solid-phase peptide synthesis to incorporate caged amino acids. Oxytocin was purchased from Tocris. Molecular weight of each compound: oxytocin (1007.2 g/mol), cOT1 (1142.3 g/mol), cOT2 (1260.4 g/mol), and cOT3 (1399.6 g/mol).

### Molecular Modeling

To understand how caging of oxytocin at different residues alter the peptide/protein interaction, molecular docking was performed for each caged oxytocin analog (cOT1, cOT2, cOT3) in comparison to oxytocin. The structure of oxytocin bound to OXTR, PDB: 7RYC was used as a template for the molecular docking experiments (Meyerowitz et al., 2022). For each designed peptide, the geometry and structure were optimized by using a fast, Dreiding-like force field via Discovery Studio (Dassault System). The generation of the receptor grid was prepared by using oxytocin bound to OXTR structure (PDB: 7RYC) and was cleaned for docking. The defined dimensions of X, Y, and Z coordinates for site-specific docking of OXTR were −140.667, 134.000, and 113.361, respectively and the box size was 86 × 126 × 118 for all three dimensions was used. Each compound was docked using Autodock Vina (Schrodinger Maestro yielded similar results). Lastly, the van-der Waals (vdW) repulsion forces between the caged oxytocin analogs and OXTR were calculated using the “Show bumps” plugin, implemented in the PyMOL interface. Heatmaps of compound binding was analyzed and plotted using Prism 8 (GraphPad, San Diego, CA).

### Photophysical and photochemical characterization

All measurements were performed in the dark or under red light. Samples were stored at –20 °C. UV-Vis spectroscopy was performed using a Cary 60 UV-Vis Spectrophotometer equipped with a PCB 1500 High-Performance Peltier thermostat (Agilent Technologies, Santa Clara, CA). Samples were measured using ultra-micro cuvettes with a 10 mm light path (BrandTech, Essex, CT). Irradiation was performed using an Optoscan Monochromator equipped with an Optosource High-Intensity Arc Lamp (Cairn Research, Kent, UK) set to 15 nm full width at half maximum and installed with a 75 W UXL-S50A xenon arc lamp (USHIO, Tokyo, JP). The light source was set to 365 nm, and the power output was measured at the sample position using a PM100D compact power and energy meter console and S120VC Si photodiode sensor (Thorlabs, Newton, NJ). A constant 3 mW power output at 365 nm was maintained throughout the experiments. The monochromator was controlled using a custom MATLAB program, and data were analyzed and plotted using Prism 8 (GraphPad, San Diego, CA).

For measuring the UV-visible absorption spectra, we prepared a 20 mM DMSO stock solution of each respective compound, diluted to 20 μM (1% DMSO in PBS). A sample (1 mL) of the solution (20 μM) was transferred to a cuvette and measured by under ambient conditions (24 °C).

For uncaging, a sample of the photocaged compound cOT1 (1 mL, 20 μM, 1% DMSO in PBS) was warmed to 37 °C and kept at this temperature throughout the irradiation period. The sample was irradiated with 365 nm monochromatic light (3 mW) for different time durations, and the corresponding absorption spectra were measured under the abovementioned conditions. Further analysis of peptide uncaging was done using mass spectrometry for cOT1 and cOT2. A sample of the respective photocaged compound (1 mL, 20 μM, 1% DMSO in PBS) was warmed to 37 °C and kept at this temperature throughout the irradiation period. The sample was irradiated with 365 nm monochromatic light (3 mW) for different durations. A sample (50 μL) was taken at each timepoint after thorough mixing then transferred to an amber vial. Samples were injected (10 μL) and characterized by LC-MS using a standard method (5–100% MeCN in H_2_O, 0.1% formic acid, 5 min runtime, 1 mL/min). The total ion count (TIC) for the proton adducts of oxytocin and of each photocaged compound were extracted from each run by integration of the extracted peak, and the area of the TIC peak for oxytocin and the respective photocage were plotted at each timepoint. The data for the TIC of each compound is shown as the average of two replicates. LC-MS traces from a representative set of measurements for each caged derivative were overlaid showing a time course of photouncaging with reduction of the photocaged compound and simultaneous increase in oxytocin. Oxytocin is stable towards 365 nm irradiation at the durations of experiments performed. The extent of ionization is compound-dependent. LC-MS Data:

OT: LC-MS t_R_ = 2.19 min (5–100% MeCN in H_2_O, 0.1% formic acid, 254 nm detection)

LRMS (ESIP) for C H N O S ^+^ [M+H]^+^: calcd.: 1007.4, found 1007.4.

cOT1: LC-MS t_R_ = 2.82 min (5–100% MeCN in H_2_O, 0.1% formic acid, 254 nm detection)

LRMS (ESIP) for C H N O S ^+^ [M+H]^+^: calcd.: 1142.48, found 1142.5.

cOT2: LC-MS t_R_ = 2.97 min (5–100% MeCN in H_2_O, 0.1% formic acid, 254 nm detection)

LRMS (ESIP) for C_53_H_76_N_13_O_18_S_2_^+^ [M+H]^+^: calcd.: 1246.5, found 1246.5.

### Cell culture

Stably transfected CHO-K1 cell lines overexpressing OXTR or V1a were obtained from Dr. Bryan Roth via the National Institute of Mental Health (NIMH) Psychoactive Drug Screening Program. The cells were cultured and maintained in Hams F12 medium (Gibco, Waltham, MA) supplemented with 400 μg/ml G418 (Gibco, Waltham, MA), 10% heat-inactivated FBS (Gibco, Waltham, MA), 15 mM HEPES (Gibco, Waltham, MA), 100 units ml^−1^ of penicillin, 100 μg ml^−1^ of streptomycin. The cells were grown in an atmosphere of 37°C and 8% CO_2_ and were passaged following trypsinization with Trypsin-LE (Gibco, Waltham, MA).

#### Calcium imaging

CHO-K1 cells overexpressing OXTR were seeded overnight at 250,000 cells/250 µL on 35 mm glass-bottom dishes (Ibidi, Fitchburg, WI) pretreated with poly-lysine. The cells were incubated with 250 µL of dye-loading solution using abcam Fluo-8 No Wash Calcium Assay kit (ab112129) for 1 hour at room temperature. Imaging was carried out on a Nikon Ti widefield - Nikon TIRF microscope. The cell samples were treated with either buffer (HBBS), oxytocin, and cOT1 to achieve a final concentration of 1 µM each. For the uncaging experiment of cOT1, LED uncaging used full-field illumination through the epi-fluorescence pathway with a 365 nm LED for 1 second (CoolLED pe-4000 illumination system) with a power output of 3 mW.

#### Calcium flux assay

CHO-K1 cells overexpressing OXTR were seeded overnight at 50,000 cells/100 µL/well in a black wall/clear bottom 96-well plate pretreated with poly-lysine (Gibco, Waltham, MA). The cells were incubated with 100 µL of dye-loading solution using an abcam (Cambridge, United Kingdom) FLUO-8 No Wash Calcium Assay kit (ab112129) for 1 hour at room temperature. Buffer (HBBS), oxytocin, and the caged oxytocin analogs (cOT1, cOT2, cOT3), either caged or uncaged (with 365 nm light), was added (25 µL/well) to achieve a concentration of 1 µM respectively. LED uncaging of the compounds was done in cuvette with a 365 nm LED (CoolLED pe-4000 illumination system, Andover, MA) for 30 minutes with a power output of 3 mW to achieve maximum photorelease of oxytocin. The calcium flux assay was carried out on a FlexStation 3 Microplate Reader (Molecular Devices, San Jose, CA).

#### Radioligand binding assay

Radioligand binding assays were carried out by the (NIMH) Psychoactive Drug Screening Program (Besnard et al., 2012). The detailed experimental protocols for the radioligand assay are available on the NIMH PDSP website at http://pdsp.med.unc.edu/UNC-CH%20Protocol%20Book.pdf. Samples were provided either caged or uncaged (with a 365 nm LED for 30 min with a power output of 3 mW). Binding data were analyzed and plotted using Prism 8 (GraphPad, San Diego, CA).

### Immunohistology

Mammary tissue was cleared (Lloyd-Lewis et al., 2016) and stained with polyclonal oxytocin receptor antibody. The mammary gland was excised and fixed overnight from either lactating wild-type (C57BL/6) or oxytocin knockout mice in 4% PFA (Electron Microscopy Sciences, Hatfield, PA). Tissue was incubated at 37 °C in CUBIC Reagent 1 A (10 wt% Triton, 5 wt% N,N,N’,N’-tetrakis (2-HP) ethylene diamine, 10 % Urea, NaCl 25 mM) clearing solution for 4 days. Samples are rinsed 3X in phosphate-buffered saline (PBS) and then incubated at 4 °C for 4 days in primary antibody for oxytocin receptor (polyclonal antibody generated in-house) at a concentration of 1μg/ml. Subsequently, the samples were diluted in PBS with Triton (PBST) containing 10% serum, rinsed again, and then incubated at 4 °C in secondary antibody (AlexaFluor goat anti-rabbit A32732, ThermoFisher, Waltham, MA) for 2 days. Lastly, the samples were rinsed 3X, then cleared in CUBIC Reagent 2 (50 w/v% Sucrose, 25 w/v% Urea, 10 w/v% Triethanolamine 0.1 w/v% Triton) at 37 °C for 24 h. The cleared and stained tissue was imaged using Zeiss LSM880 confocal microscope.

### *Ex vivo* mammary tissue imaging

Mammary glands were harvested from lactating GCaMP6f;K5CreERT2 mice diced into 3- to 4-mm^3^ pieces and loaded with CellTracker Red (1.5 μM, ThermoFisher C34552, ThermoFisher, Waltham, MA) for at least 30 min in DMEM/F12 containing 10% FCS. Under these conditions, CellTracker preferentially labels luminal cells (Stevenson et al., 2020). Tissue was bathed in 85 nM cOT1 for at least 5 min before imaging. Samples were imaged on a Leica TCS SP8 inverted microscope (Leica, Germany) with 20x 0.75NA HC PL APO CS2 objective. For uncaging, a region of interest (ROI) was targeted with a 405 nm laser at 100% laser power for 3s each trial. A final concentration of 85 nM of free oxytocin was added in bath for oxytocin wash. Protocol for image processing and analysis was previously reported in (Stevenson et al., 2020).

It is worth noting the uncaging in this experiment was done with a 405 nm laser at the highest power, which is not optimized (less than 10% efficiency) for uncaging the oNBZ cage group of cOT1 (Klán et al., 2012). We are confident that using a UV laser with a wavelength between 340 – 385 nm would significantly improve uncaging efficiency. However, it is noteworthy that succeeding in uncaging cOT1 with 405 nm light broadens the access and utility of this tool because most standard microscope setups are equipped with 405 nm lasers. Additionally, this experiment demonstrates the potential for experiments where three wavelengths multiplexing could be useful. Here we uncaged oxytocin with UV light and tracked the cells in the red channel and calcium imaged in the green channel.

### *In vitro* electrophysiology

#### CA2 Hippocampus

*In vitro* recordings were performed in acute slices containing dorsal hippocampus prepared from male and female, 2-5 month old, OXTR-cre::Ai9 mice. Mice were anesthetized with a mixture of ketamine/xylazine (150 mg/kg and 10 mg/kg, respectively) and perfused transcardially with an ice-cold sucrose solution containing (in mM): 206 Sucrose, 11 D-Glucose, 2.5 KCl, 1 NaH_2_PO_4_, 10 MgCl_2_, 2 CaCl_2_ and 26 NaHCO_3_, bubbled with 95% O_2_-5% CO_2_. Following perfusion and decapitation, brains were removed and placed in the cold sucrose for sectioning using a Leica VT 1200S Vibratome (Leica, Germany). Transverse, 300 μm sections of left and right hippocampus were cut and transferred to an oxygenated, 34°C recovery chamber filled with artificial cerebro-spinal fluid (ACSF) containing (in mM): 122 NaCl, 3 KCl, 10 D-Glucose, 1.25 NaH_2_PO_4_, 2 CaCl_2_, 2 MgCl_2_, and 26 NaHCO_3_, bubbled with 95% O_2_-5% CO_2_. After incubation, slices were held in bubbled ACSF at room temperature for up to 6 h until recordings were made.

For recording, slices were placed in a submerged slice chamber continuously perfused with ACSF at a rate of 1–3 mL/min and maintained at a bath temperature of 30°C. tdTomato-positive neurons in the CA2 pyramidal cell layer were visualized with LED illumination under an upright microscope. Whole-cell patch-clamp recordings were performed as described previously (Tirko et al., 2018), using a MultiClamp 700B amplifier (Axon Instruments, Union City, CA) and pCLAMP version 10.7.0.2 (Molecular Devices, San Jose, CA) for data collection. Signals were filtered at 10 kHz and sampled at 20–50 kHz with a Digidata 1440 data acquisition interface. During recordings, patch pipettes with a resistance of 3~5 MΩ were made from borosilicate glass (World Precision Instruments, Sarasota, FL) with a Sutter Instrument P-97 micropipette puller (Novato, CA) and filled with a solution containing (in mM): 126 K-gluconate, 4 KCl, 10 HEPES, 4 Mg-ATP, 0.3 Na_2_-GTP, 10 phosphocreatine (pH to 7.2 with KOH).

To assess effects of oxytocin, a stable baseline was established for 5 min and 1 μM oxytocin (Tocris) in ACSF was washed on for 30 min. To assess effects of cOT1 uncaging, after baseline stability was achieved in ACSF, 1 μM cOT1 was washed in for another 5-10 min to maintain the stable baseline. Then a CoolLED pe-4000 illumination system was used to generate 365nm of UV light pulsed at 20Hz for 1 to 2 min over the slices through the objective. For further control experiment using OXTR antagonist OTA, OTA was continuously presented before the wash-in of cOT1 to the end of the recording. The whole set of experiments was conducted in dark as much as possible to avoid pre-emptive uncaging due to ambient UV light. Peak depolarization was measured as the mean membrane potential over a 2-minute period of highest membrane depolarization at least 10 minutes after uncaging of cOT1, subtracted from the mean membrane potential before the light stimulation. Input resistance and series resistance were monitored throughout the experiments, and recordings were rejected if series resistance increased to above 25MΩ or more than 20%.

#### Experimental design and statistical analysis

Experiments were not performed blindly. In all cases, four or more animals with both sexes were used for each parameter collected and were pooled for analysis. Each recorded neuron came from one brain slice of one experimental animal. There was no repeated use of any brain slice. Individual sample sizes for slice patch clamp recording (n=number of neurons, included in each figure legend) are reported separately for each experiment. All statistical analysis was performed using GraphPad Prism 9. Statistical comparisons before and after the uncaging of cOT1 were made using paired two-tailed Student’s t-test. Statistical comparisons for different groups were made using one-way or two-way ANOVA and post hoc Turkey’s test. Each statistical method is clearly stated in the result section or the figure legends. All statistical tests were two sided. Data distribution was assumed to be normal, but this was not formally tested. Data are presented as mean ± sem. Individual data points are plotted in figures. All raw data sets are openly accessible upon request.

#### Auditory Cortex

*In vitro* recordings were performed in acute slices of the auditory cortex prepared from male and female 2-5 month old OXTRcre::Ai9 mice. Mice were deeply anesthetized with 5% isoflurane and perfused transcardially with ice-cold dissection buffer containing (in mM): 87 NaCl, 75 sucrose, 2.5 KCl, 1.25 NaH2PO4, 0.5 CaCl2, 7 MgCl2, 25 NaHCO3, 1.3 ascorbic acid, and 10 D-glucose, bubbled with 95%/5% O2/CO2 (pH 7.4). Following perfusion and decapitation, brains were removed and placed in the cold buffer solution, and slices (250–300 μm thick) were prepared with a vibratome (Leica P-1000). Slices were placed in warm artificial cerebrospinal fluid (ACSF, in mM: 124 NaCl, 2.5 KCl, 1.5 MgSO4, 1.25 NaH2PO4, 2.5 CaCl2, and 26 NaHCO3) (33–35 °C) for <30 min, then cooled to room temperature (22–24 °C) for at least 30 min before use. Slices were transferred to the recording chamber and superfused (2.5–3 ml min−1) with oxygenated ACSF at 33 °C.

Whole-cell current-clamp recordings were made from layer 2/3 pyramidal cells with a Multiclamp 200B or 700B amplifier (Molecular Devices, San Diego). Cortical regions and layers were identified using IR video microscopy (Olympus, Japan), and OXTR-positive neurons were identified based on expression of tdTomato, visualized with an Olympus 40 × water-immersion objective with TRITC filter. Data were filtered at 2 kHz, digitized at 10 kHz, and acquired with Clampex 10.7 (Molecular Devices, San Diego, CA).

To assess the modulatory effects of oxytocin and caged oxytocin (cOT1), whole-cell recordings were acquired in current-clamp configuration with patch pipettes (4–8 MΩ) containing the following intracellular solution (in mM): 127 K-gluconate, 8 KCl, 10 phosphocreatine, 10 HEPES, 4 MgATP, 0.3 NaGTP, pH 7.2. To assess the effects of oxytocin, a stable baseline was established for 3–10 min, and 1 μM oxytocin (Tocris, United Kingdom) in ACSF was washed on for 10–15 min, followed by a washout period. To assess the effects of cOT1, after baseline stability was achieved and following 5-10 minutes of cOT1 wash-in, 365 nm light was pulsed at 20-Hz for 1s over the recording chamber using a CoolLED pe-4000 illumination system. The peptide (cOT1) was pre-emptively kept in the dark as much as possible to avoid uncaging due to ambient UV light. Peak depolarization was measured as the mean membrane potential over the 2-minute period of highest membrane depolarization at least 10 minutes after wash-in of cOT1, subtracted from the mean membrane potential during the 2 minutes of the baseline period immediately before initiation of drug wash-in.

#### Statistical analysis

Current clamp recordings were analyzed offline using Clampfit 10.7 (Molecular Devices, San Diego, CA). Recordings were excluded from analysis if the access resistance (Ra) changed >30% compared to baseline. Spontaneous PSCs were detected offline using pClamp software for event detection. Student’s t-test was used to compare two groups using Prism 9.0 GraphPad software. Data is displayed as the mean +/− the standard error of the mean (s.e.m.).

### Behavior

Pup-naïve female C57BL/6 wild-type virgin mice were bred and raised at NYU Grossman School of Medicine and kept isolated from both dams and pups until approximately five weeks old. Before implantation with an optofluidic cannula (details below), naïve virgins were prescreened for pup retrieval (detailed below) to exclude spontaneous retrievers or pup mauling. Typically, <30% of naïve virgins retrieve at least one pup or maul pups during pre-screening (Marlin et al. 2015). Dams were also pre-screened to ensure they retrieved pups before using them in this study.

#### Optofluidic cannula implantation

In brief, a total of 30 female C57BL/6 wild-type mice at 5 weeks of age were anesthetized and implanted with 1 mm Optical fiber Multiple Fluid Injection Cannula system (Doric Lenses, Canada) (part: OmFC_SM3_400/430-0.66_1.0mm_FLT_0.5mm) targeting the auditory cortical region—coordinates: AP: −2.8 mm and ML: −4.35 mm. The implant was attached to the skull with dental cement. Post-surgery the mice were group-housed (cannula was protected using a cap) for 4 - 5 weeks to allow recovery from the surgery. Daily food intake and body weights were monitored. Then each animal was screened again for spontaneous pup retrieval for the baseline experiment. 7 animals were removed during recovery due to bodyweight drop (1 animal) or spontaneous pup retrieval post-surgery (6 animals).

Post hoc verification of implantation was conducted after the completion of behavior tests. Animals were sacrificed and fixed with 4% paraformaldehyde via transcardial perfusion. Then, 100 μm brain slices were cut using a cryostat, and cannula and optic fiber tracks were viewed under a stereo microscope. Confocal images were taken using Zeiss LSM700 or LSM800 (Zeiss, White Plains, NY). Data from mice with incorrect cannula placement were excluded from the data analysis (2 animals).

#### Pup retrieval

A single test session consists of 10 individual trials of pup retrieval for an individual mouse. The pup retrieval test was performed as pre-screening before surgery, baseline, and post-infusion with caged oxytocin (cOT1). The mice were placed in a behavioral arena (38 × 30 × 15 cm) for prescreening or a customized behavioral arena setup of identical size for the baseline and post-infusion sessions. The mice were first given 20 minutes to acclimate to the behavioral arena before each testing session. Subsequently, at least 4 pups ranging from postnatal day 1–4 was placed in a corner of the arena under nesting material. A single pup was removed from the nest and placed in an opposite corner of the arena for each trial. The experimental mouse was given ten two-minute trials to retrieve the displaced pup back to the nest. If the displaced pup was not retrieved within two minutes, the trial was scored as a failure, and the pup was returned to the nest. If the pup was retrieved back to the nest, the time of retrieval was recorded. This is repeated until ten trials are complete but each time removing a different pup from the nest. We consider reliable retrieval when the mouse successfully retrieves pups in at least two out of ten trials. After ten trials, the pups were returned to their home cage with their dam. An ultrasonic microphone (Avisoft, Germany) was used during all pup retrieval experiments to confirm that the isolated pups vocalized distress calls.

After the baseline test for pup retrieval was performed, each virgin mouse was cohoused with a dam and her pups. After 1 hour, the virgin mouse is placed back into the customized behavioral setup equipped for delivering fluid containing cOT1 and 365 nm light for uncaging using both a fluid microinjector (World Precision Instruments, United Kingdom) and an LED (Thor Labs 365 nm LED controlled by High-Power 1-Channel LED Driver with Pulse Modulation, Newton, NJ) respectively. All the female virgin mice were infused with cOT1 (50 μM in saline, 1.5 μl at 1 μl/min). Immediately after the infusion was complete, one group (n = 9) of the mice had the 365 nm LED light turned on (385 nm, 10 mW, 500 ms, 2 Hz) for the photorelease of native oxytocin and the other group (n= 12) had it off. The mice were then untethered from the LED and microinjector and allowed to complete the 20-minute acclimation period to the behavior arena before pup retrieval testing. Then the 10 trials of pup retrieval were carried out as described previously (Marlin et al., 2015). Pup retrieval was tested after infusion of cOT1 with or without light for the following time points: 1, 3, 6, 12, 18, 24, 36, 48, and 72 hours. Fisher’s two-tailed exact test was used to compare the LED on group (uncaged) versus the LED off group (caged).

## Acknowledgements

We thank Michael Cammer for assistance with microscopy and Dr. Bryan L. Roth for providing us with OXTR and AVRP CHO-1 Cells. I.A.A is supported by the NIH (5T K00 MH123667) and the Burroughs Wellcome Fund PDEP award. C.J.B.M acknowledges the support of the Natural Sciences and Engineering Research Council of Canada (NSERC). B.E.H. and C.J.A. are supported by the NYU MacCracken Fellowship. M.V.C. acknowledges the support of the NIH (R01 MH119136-04 and U19 NS107616-04). F.M.D. is supported by the National Health and Medical Research Council (NHMRC) of Australia (1138214, 1141008 and 2003832) and a Young Investigator Award from the Novo Nordisk Foundation (NNF20OC009705). R.C.F. and R.W.T. are supported by the NIH BRAIN Initiative (NS107616), and R.C.F. is supported by the NICHD (HD088411).

## Contributions

I.A.A. designed and synthesized peptides and carried out molecular simulations. B.E.H. and C.J.A. performed photochemical and photophysical characterization of peptides. I.A.A. performed cell culture experiments. L.K. and P.C. performed clearing tissue imaging of mammary tissue. K.A.G. and F.M.D. performed mammary imaging and G.C.V. performed analysis. J.J.L. and C.J.B.M performed electrophysiology in brain slices. J.J.L. performed cannula implant surgeries. I.A.A., A.B.A., and A.T.S. performed behavior experiments. I.A.A., F.M.D., J.J.L., and C.J.B.M made figures. I.A.A. and R.C.F. conceived and managed the project. I.A.A., M.V.C., F.M.D., D.T., and R.W.T. provided technical expertise and scientific direction. I.A.A. and R.C.F. wrote the manuscript.

## Ethics Declarations

Competing interests

The authors declare no competing interests.

